# A Data-driven Individual-based Model of Infectious Disease in Livestock Operation: A Validation Study for Paratuberculosis

**DOI:** 10.1101/394569

**Authors:** Mohammad. A. Al-Mamun, Rebecca. L. Smith, Annette. Nigsch, Ynte. H. Schukken, Yrjo.T. Gröhn

## Abstract

Chronic livestock diseases cause large financial loss and affect the animal health and welfare. Controlling these diseases mostly requires precise information on both individual animal and population dynamics to inform farmer’s decision. Mathematical models provide opportunities to test different control and elimination options rather implementing them in real herds, but these models require valid parameter estimation and validation. Fitting these models to data is a difficult task due to heterogeneities in livestock processes. In this paper, we develop an infectious disease modeling framework for a livestock disease (paratuberculosis) that is caused by *Mycobacterium avium* subsp. *paratuberculosis* (MAP). Infection with MAP leads to reduced milk production, pregnancy rates, and slaughter value and increased culling rates in cattle and causes significant economic losses to the dairy industry in the US. These economic effects are particularly important motivations in the control and elimination of MAP. In this framework, an individual-based model (IBM) of a dairy herd was built and a MAP infection was integrated on top of it. Once the model produced realistic dynamics of MAP infection, we implemented an evaluation method by fitting it to data from three dairy herds from the Northeast region of the US. The model fitting exercises used least-squares and parameter space searching methods to obtain the best-fitted values of selected parameters. The best set of parameters were used to model the effect of interventions. The results show that the presented model can complement real herd statistics where the intervention strategies suggested a reduction in MAP but no elimination was observed. Overall, this research not only provides a complete model for MAP infection dynamics in a cattle herd, but also offers a method for estimating parameter by fitting IBM models.

## Introduction

Chronic livestock diseases like paratuberculosis (PTB) and bovine tuberculosis (bTB) are commonly reported worldwide (1,2). Bovine TB is caused by the pathogen Mycobacterium bovis (*M. bovis*) while PTB is caused by *Mycobacterium avium* subsp. *paratuberculosis* (MAP). In the UK, bTB has been spreading over the last two decades, putatively due to the presence of a wildlife reservoir in badgers(3). In United States (US), 68% of dairy herds have apparently at least one cow that is infected with MAP (4). Both diseases pose a potential threat not only to animal health and production, but also to public health. Historically, bTB has been a contributor to human TB cases worldwide and PTB infections in humans have been associated with an increased risk of Crohn’s disease in humans(5). Recently, it has been reported that these diseases may induce additional collateral risks for public health due to dispensed antibiotics as a treatment in some cases can potentially contribute to the spread of antibiotic resistance(6).

In the US cattle industry, the cost of PTB was estimated at $250 million every year (7). Infection by MAP usually occurs in the first year of life(8) and transmission can occur vertically (9) and/or horizontally via ingestion of fecal material contaminated by MAP (10). As PTB is a slowly progressive disease, progression of individual animals through different MAP infection states is a complex continuous process alternating excreting and non-excreting stages with a late onset of clinical signs (11,12). It has a large economic impact for producers due to decreased milk production (13–15), premature culling (16,17), reduced slaughter value (18), low fertility (19,20), and an increased animal replacement rate (21). However, tests routinely used on individuals have low sensitivities, especially in the early stages of the disease (22).

In last two decades, different mathematical models have been developed on a within-herd scale to understand MAP transmission dynamics (23,24) and effectiveness of recommended control strategies (25–28). These models were simulated to assess the impact of contact structure on the MAP transmission (23), efficacy of test-and-cull policy (24,25,29,30), impact of low diagnostic test sensitivity in decision making (8,31), stopping some transmission pathways using hygiene improvement (32), improved calf management (33), impact of super-shedders in transmission(34,35), and economic efficacy of recommended programs (29). Most of these studies suggest that culling a test positive animal is an effective solution to reduce the prevalence. However, none of the previous models considered the pervasiveness of MAP in the farm environment and the value of information of individual animal along with real dairy herd data. Moreover, controlling MAP requires significant management of testing and culling strategies to reduce the prevalence, which are normally unregulated and reliant on farmers’ decisions(36). The decision of culling an animal is not straightforward and poses a multiscale problem where an individual animal, farm dynamics, infectious status and disease symptoms, and management profit are related (37). Substantial costs are also related to the implementation of control measures and prevention (21,34,38). Though previous compartmental MAP models have shown many potential interventions programs, most considered population-level decision making rather than individual-level animal information. Recently, individual-based models (IBMs) have been proposed to show the value of the information about the infection, daily life events and management policy for each individual animal within the farm (32,37,39–42)

Mathematical models of infectious diseases often direct us to understand both infection biology and efficacy of intervention policies taken in human and veterinary medicine(43). However, translating modeling results into practice require appropriately real-world assumptions to be built into the model. We hypothesize that in case of MAP, the use of model results will more realistic when the model has been built on up-to-date infection biology and epidemiology, parametrized from adequate real herd data and fitted back to that real-world scenario to test the recommended intervention strategies. In this paper, our aim is to build an IBM framework of MAP infection that is fitted to and validated by in-depth longitudinal data from three northeastern dairy farms. The objective of this study was four-fold: first, we extended an existing IBM of a dairy herd to resemble the population level parameters (i.e. milk yield, herd size) with three real herds to create three *in silico* herds; second, we fitted the milk-yield measurement of individual animal to those herds; third, we fitted the model-predicted apparent MAP prevalence to the observed data to obtain herd-specific important infection parameters; and fourth, we integrated a risk-based control strategies on those three *in silico* herds to evaluate the efficacy. Finally, we discuss the value of observational data to feed information to simulation models, thereby making simulations more reflective and predictive of real-world circumstances.

## Materials and Method

### The Individual-based model

We used the dairy herd model named a multiscale agent-based simulation of a dairy herd (MABSdairy), an improved version of dairy herd published in Al-Mamun et al. (32,40). The MABSdairy is a multiscale stochastic IBM that simulates individual cows in a standard US cattle herd with a daily time step. In brief, each cow resides in one of three different management operations: adult/milking (aged >720 days), calf (aged 1-60 days) and heifer rearing housing (aged 61-719 days). Adult cows must calve to produce milk and the lactation cycle refers to the period between one calving and the next. The lactation cycle included the processes of a voluntary waiting period (interval during the postpartum period), insemination, and the dry off period (a non-lactating period prior to an impending parturition to optimize milk production in the subsequent lactation). For the fitting purpose, we modified the milk production Wood lactation curve by adding a herd-specific term and a herd-specific random component(44). The function is defined as

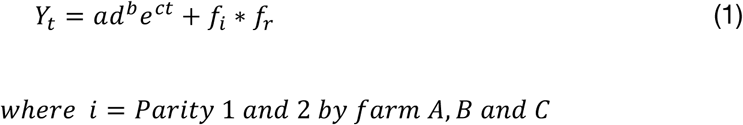

where *Y*_*t*_ is the yield on day t after calving, d is days in milk (DIM), *a* is a scaling factor for initial yield, *b* is a rate factor for the increase in yield to peak, *c* is a rate factor for the decline after the peak, f_i_ farm specific factor and f_r_ is a random number. We used base milk yield parameters from Dematawewa et al. for parities 1 and ≥2 in the basic model (45).

### MAP infection dynamics

The infection compartments in the milking herd were divided into four categories: susceptible (X_A_), latent (H), low shedding (Y_1_), and high shedding (Y_2_). In calf rearing housing, there were two infection categories: susceptible (X_C_) and infected (Y_C_). In heifer rearing housing, there were also two infection categories: susceptible (X_H_) and infected (Y_H_). We included six different transmission routes: adult-to-adult, adult-to-calf (vertical transmission), adult-to-calf (horizontal transmission), environmental contamination, calf-to-calf, and heifer-to-heifer. The detailed infection structure is shown in Fig1.

**Fig 1.**
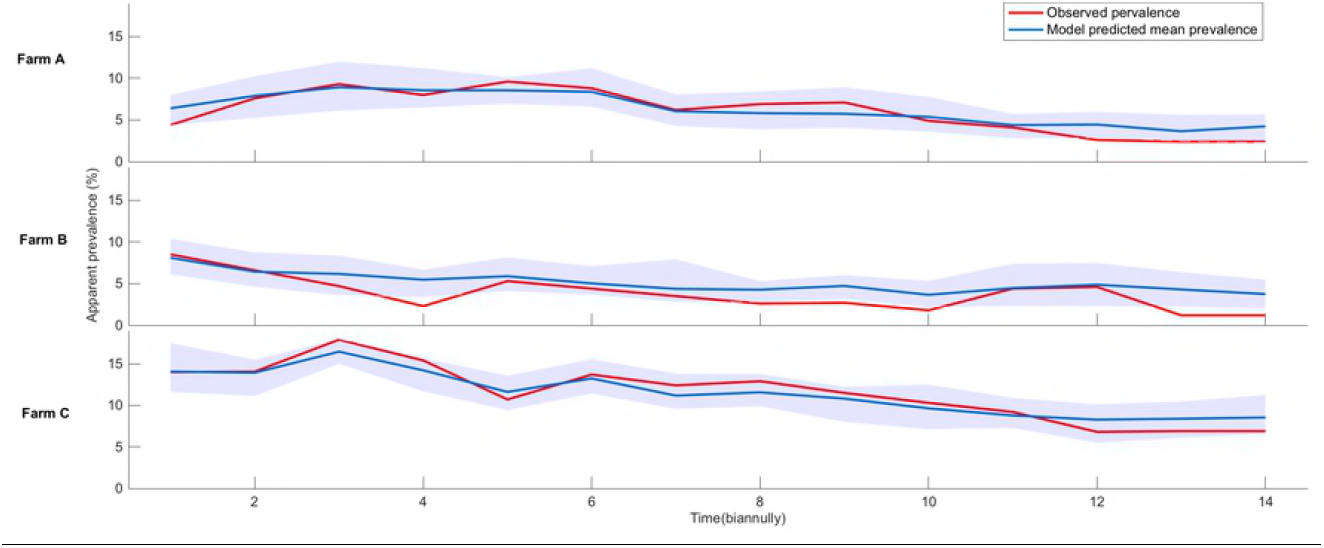
A flow diagram of animal movement among infection categories for the adult, calves, and heifers within the herd. Each horizontal gray box classifies the animals according to their initial age group. The green and red boxes define the susceptible and infected states, respectively, for each animal in the three age categories. The probabilities of exit at each time point from susceptible to latent, latent to low shedding and low shedding to high shedding animals are s_1_, h_1_, and y_1_, respectively. Vertical transmission probabilities from latent, low shedding and high shedding animals are V_h_, V_y1_, and V_y2_, respectively. Horizontal transmission probabilities to calves from low shedding and high shedding animals are H_y1_ and H_y2_, respectively. The probability an animal gets infected by the environment is *β*_*environment*_. Calf-to-calf and heifer-to-heifer transmission probabilities are C_*inf*_ and *Y*_*inf*_, respectively. Stochastic death/sale probabilities for adult, calves, and heifers are µ_a_, µ_c_, and µ_h_, respectively. µ is the replacement animals coming from heifer compartment upon completion of two years.

In the milking herd group, adult animals could be infected by low and high shedding adults. The probability of fecal-oral transmission for adult animals can be given by:

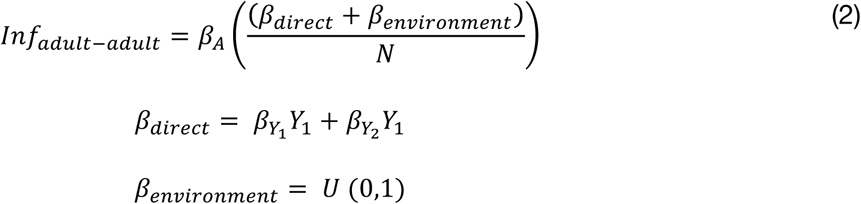

Susceptible adult animals in the milking herd compartment were susceptible to MAP infection by contact with low shedding (*Y*_*1*_) and high shedding (*Y*_*2*_) animals with transmission rates of β_Y1_ and β_Y2_, respectively. β_A_ is the adult to adult transmission coefficient, β_*environment*_ is the MAP contamination risk from the environment and *N* is the total number of animals in the milking herd, *N= X*_*A*_*+H+Y*_*1*_+*Y*_*2*_. The horizontal infection probability to calves can be determined by

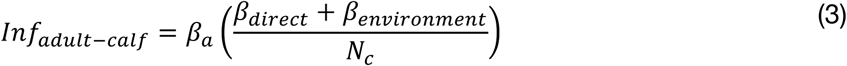

*β*_*a*_ is the horizontal transmission coefficient for an adult to newborn calves and N_c_ is the total number of calves at every day, N_c_ = X_c_ + Y_c_. A calf can also become infected vertically (i.e., in utero infection) by an adult and which are modelled using the certain proportions(25).

A calf stays in calf rearing housing for the first 60 days after birth. The probability of direct transmission was calculated as

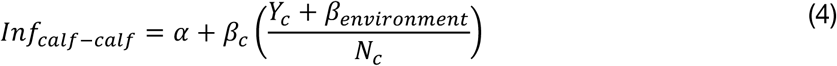

*β*_c_ is the horizontal calf-to-calf transmission coefficient, N_c_ is the total number of calves at each day, *X*_*c*_ is susceptible calves, *Y*_*c*_ is infected calves. During the first day after birth, a calf may also be infected horizontally by infected adults present in the maternity pen or vertically by an infected dam.

Susceptible calves became susceptible heifers and infected calves became infected heifers. Infected heifers could infect susceptible heifers by the heifer-to-heifer transmission path

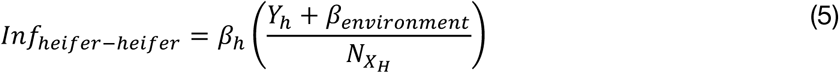

β_*h*_ is the horizontal heifer-to-heifer transmission coefficient, and the total number of heifers is N_XH_ = X_H_ + Y_H_. After one year, the infected heifers became latent heifers and eventually entered the milking herd as latent adults. For simplifying the model, we assumed that heifer remains in the heifer rereading housing are transiently shedding while they ended up in the adult herd as latent animals.

### Observed herd data

The longitudinal dataset used here was obtained from a longitudinal study of three commercial dairy farms in the northeastern US: farm A in New York State, farm B in Pennsylvania, and farm C in Vermont (46,47). All three farms participated in the Regional Dairy Quality Management Alliance (RDQMA) project, which was a multistate research program conducted under a cooperative research agreement between the USDA Agricultural Research Service (ARS) and four Universities: Cornell University, Pennsylvania State University, University of Pennsylvania, and University of Vermont. The project consisted of longitudinal data collection for endemic infectious diseases of public and animal health concern in dairy herds. For a more complete description, including information on farms, samplings, and microbial analyses, see Pradhan et al.(46) Briefly, the milking herds consisted of approximately 330, 100, and 140 cows on farms A, B, and C, respectively. Sampling commenced in February, March, and November 2004 on farms A, B, and C, respectively, and continued for approximately 7 years, until 2010. The project design included a biannual collection of individual fecal samples and a quarterly collection of individual serum samples from all milking and non-lactating cows. Additionally, culled cows were tracked as much as possible from the farm to the slaughterhouse, where four gastrointestinal (GI) tissues and a fecal sample were collected with the cooperation of USDA Food Safety and Inspection Service (FSIS) personnel. The harvested tissues included two lymph nodes located at the ileocecal junction and two pieces of ileum, one taken from 20 cm proximal to the ileocecal valve and the other taken from very near the ileocecal valve. In addition to the sampling of animals, the farm environment was sampled in approximately 20 locations on a biannual basis. All fecal and environmental samples were tested by 4-tube culture for presence of viable MAP organisms, reported as colony-forming units per tube. All serum samples were tested using the ParaCheck ELISA (Prionics USA Inc., La Vista, NE) for antibody reactions to MAP antigens. On each of the farms, demographic data, production data and herd management information was collected. Precise demographic data included birth date, birth location, calving dates, fertility data, animal location data (pen status at any point in time), dry-off dates and culling information and cull dates. These demographic data were collected for each animal present on the farms. All infection data, strain typing data, herd management, demographic and production data was maintained in a relational database.

### Model parameters

The parameterization of the base dairy herd model is described in Al-Mamun et al. (32,48). Initial infection parameter values were updated according to Mitchell et al. 2015 (43). Table 1 provides the base parameters for the initial MAP transmission before fitting the model to the RDQMA herds.

**Table 1.** Base parameter values of *Mycobacterium avium* subsp. *paratuberculosis* (MAP) infection within a dairy herd.

| Symbols | Description | Initial value | References |
| --- | --- | --- | --- |
| $V_h$ | The proportion of calves from latent animals infected at birth | 0.15 | (25) |
| $V_{y1}$ | The proportion of calves from low-shedding animals infected at birth | 0.15 | (25) |
| $V_{y2}$ | The proportion of calves from high-shedding animals infected at birth | 0.17 | (25) |
| $\beta_A$ | Adult-to-adult transmission coefficient | 0.05 | (32) |
| $\beta_a$ | Adult-to-calf transmission coefficient | 0.383 | (32) |
| $\beta_c$ | Calf-to-calf transmission coefficient | 0.0025 | (32) |
| $\beta_h$ | Heifer-to-heifer transmission coefficient | 0.001 | Calibrated in the model |
| $\beta_{y1}$ | Transmission rate between low shedders ($Y_1$) and susceptible ($X_A$) | 2/year | Calibrated in the model |
| $\beta_{y2}$ | Transmission rate between high shedders ($Y_2$) and susceptible ($X_A$) | 20/year | Calibrated in the model |

### Model fitting method

The goal of the model-fitting exercise was to estimate key parameters in order to produce results consistent with the epidemiologic data from three farms. Our fitting exercise was two-fold: first, we fitted our base dairy herd models with farm-specific parameters (total population and milk yield), then we fitted the model predicted apparent prevalence results based on antemortem ELISA and fecal testing and postmortem tissue and fecal testing results for the farm. To assess the goodness-of-fit we sampled from the defined parameter ranges in multiple rounds and the model was run for three different scenarios of each of the three farms. The model fitting was done using a non-linear fitting method named Nelder-Mead Simplex Method (49), which is used for unconstrained optimization. While fitting the milk yield and apparent prevalence, the best-fit parameters were extracted.

To determine the specific range for each parameter, we used multidimensional parameter space searching method. The point estimate of each parameter was taken as a mean value and, using Latin Hypercube Sampling, 100,000 parameter combinations were generated spanning the specified range ±75% of the mean values. The searching was done in two stages. In the first stage, we set a broad range to identify the particular regions of the parameter range and chose the best 10000 (1%) parameter sets. In the next stage, we ran the simulation with 10% parameter sets to compare with the best fit curve by minimizing the sumsquare error. The parameter ranges presented in the results section were calculated from the top 1% simulations.

### Intervention strategies

Once the three *in silico* herds were stable using fitted values, we tested a proposed intervention strategy. We chose risk-based testing and culling strategies suggested by Al-Mamun et al. (32). In brief, all cows that tested negative throughout testing were marked as low risk or green cows. The cows that tested positive were divided into two groups: yellow and red. Red animals had at least 2 positive tests out of the last 4 tests and yellow cows had one positive test. We proposed two controls: control I, culling red animals straightway (aggressive culling); and control II, culling only red animal with a delay of 305 DIM (delayed culling). The simulations results were then compared against the observed pre-fitted data from the three herds. To evaluate the efficacy of the intervention, we divided our seven years of observation into two parts: the first 4 years were used for pre-intervention fit, while the last 3 years were used for validation against the model results, in which the intervention was introduced and run for 3 years. Moreover, we extended the intervention for more two years to see the long-term efficacy.

### Simulation background

First, the base dairy herd model was initiated with a certain proportion of adult animals for farms A (330), B (100) and C (140). Second, after a 2-year burn-in period the model was run for 7 more years to resemble the observations of the real herds. During the 2 years burn-in period, each farm was assumed to be self-sufficient in producing their own replacement, so that no animal purchase from outside was needed. The model was initiated with a pre-determined distribution of animals with different parities. Every day, the algorithm first determined the group of animals. If it found adult animals, it checked reproductive status (voluntary waiting period (VWP), waiting to be inseminated, and pregnant) and milk yield status. Any cow on the 280th day of pregnancy was assumed to calve. For a newborn calf, the stillbirth probability was checked; if the calf was not stillborn, it was flagged as a calf. Only female calves were kept in the herd, and male calves were removed/sold immediately after birth. Once an adult animal calved, it transitioned to VWP status and continued in the milking herd loop until it was removed due to culling or death. Mortality was allowed in the calf rearing loop; otherwise, calves were transferred into the heifer loop at the 61st day of age. In the heifer loop, heifers were inseminated at the 400th day of age in order to become pregnant, so that they would calve at the 680th day of age. When heifers were ready to calve for the first time, they transitioned to the milking herd in the model. The model was fitted for the 7 years data for each farm. Third, for testing intervention strategies, each model was fitted to the first 4 years of data-that is called pre-intervention fit, and then the intervention was tested in 2 phases. In the first phase, 3 years and then extended more 2 years to see how the suggested strategies result in long term. The base model was developed as custom codes in MATLAB and other data analysis were done using R.

### Results

The purpose of the fitting exercises was to obtain a better fit to the estimates of three herds prior fitting to the apparent prevalence. The model predicted total number of animals (adult, calves, and heifers) closely resembles the data from the three real farms (shown in table 2).

**Table 2.** The comparison of observed and predicted values from three *in silico* farms in terms of a total number of animals, and average daily milk yield (in kg) for 305 days, presented as Mean (95% Confidence Interval).

| Herd A | Total number of animals | Milk yield: parity 1 | Milk yield: parity $\geq 2$ |
| --- | --- | --- | --- |
| Observed | 720 (708-754) | 36.07 (29.61-40.73) | 39.48 (27.11-49.86) |
| Predicted | 714 (693-737) | 36.15 (30.44-40.73) | 39.49 (27.56- 50.18) |
| Herd B |  |  |  |
| Observed | 194 (102-230) | 33.38 (24.34-40.35) | 34.52 (17.42-47.40) |
| Predicted | 200 (182-219) | 32.97 (26.51-38.31) | 34.97 (21.86-48.29) |
| Herd C |  |  |  |
| Observed | 262 (116-339) | 27.49 (19.03-34.68) | 27.49 (19.03-34.68) |
| Predicted | 221 (184-257) | 27.16 ( 20.23-32.98) | 27.90 (17.12-38.06) |

Fig2 shows the concordance between predicted and observed milk yield data from three herds. It is evident that the models predicted milk yield estimations matched with the observed milk yield from three northeastern herds. The best fit model predictions to the observed milk yield curve for parity 1 and parity ≥2 are shown in supplementary FigS1. The best fitting lines also describe that the model was able to capture inherent randomness from the data into the model. The estimation of the critical parameters *a, b, c,* and *f*_*i*_ of the modified lactation curve are presented in table 3.

**Fig 2.**
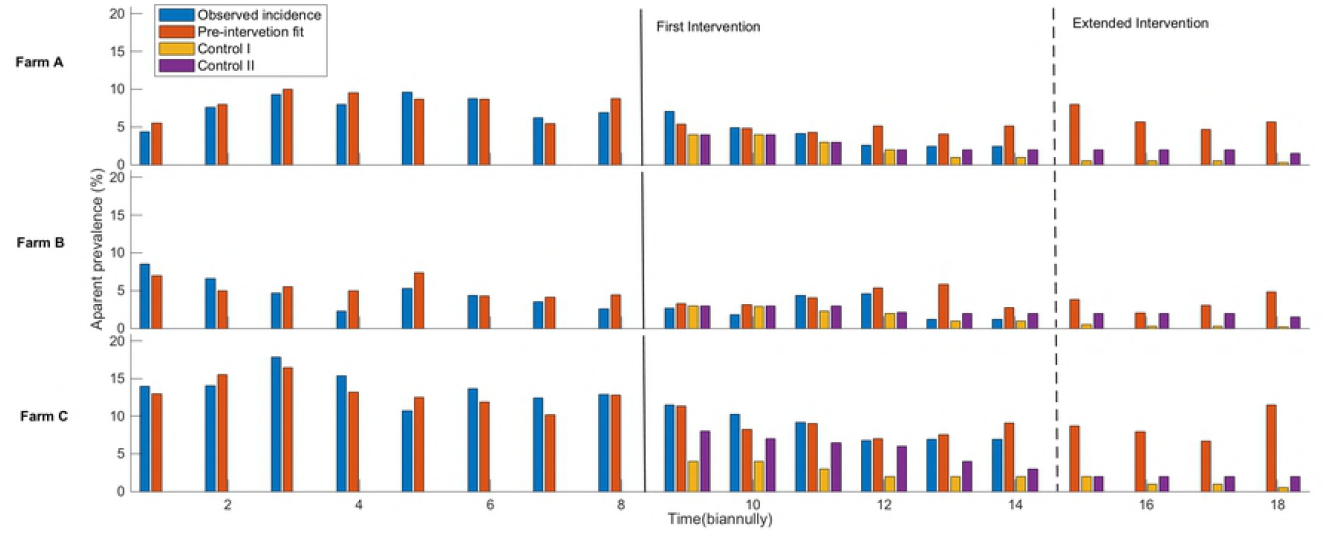
The comparison of observed and model predicted milk yield distribution for 1% simulation using best fit parameters for the milk yield. In the box plot, the bottom and top end of the bars are minimum and maximum values respectively, the top of the box is the 75th percentile, the bottom of the box is the 25th percentile, and the horizontal line within the box is median; outliers are presented as a solid black circle and the density of the milk yield is presented by the width of the violin.

**Table 3.** The estimated parameters from the fitting exercise for the modified milk yield function for three farms A, B, and C.

| | $a$ | $b$ | $c$ | Herd specific parameter<br>($f_i$) |
| --- | --- | --- | --- | --- |
| Farm A-Parity 1 | 17.87 | 0.207 | 0.00199 | 3.59 |
| Farm A-Parity$\geq 2$ | 25.23 | 0.199 | 0.00329 | 5.19 |
| Farm B-Parity 1 | 16.09 | 0.198 | 0.00196 | 6.60 |
| Farm B-Parity≥2 | 23.38 | 0.200 | 0.00392 | 8.33 |
| Farm C-Parity 1 | 15.25 | 0.193 | 0.00269 | 6.55 |
| Farm C-Parity≥2 | 16.50 | 0.215 | 0.00399 | 9.00 |

### Model fitting exercises

Table 4 represents the observed apparent prevalence and apparent incidence and the tracking of the animals in the next biannual testing for three farms for seven years, 2004-2010. The observed prevalence shows zero infected animals in the last half of 2010, for the sake of persistence scenario we replace that with the previous quarter value. During our simulation, we normalized the prevalence with the previous half of the year so that it remains consistent for our simulation. We simulated the three *in silico* farms to fit with the observed apparent prevalence data from herd A, B and C.

**Table 4.** The calculation of apparent prevalence and apparent incidence and the tracking of the animals in the next testing in bi-annually phase for three farms (2004-2010).

| Year | 2004 |  | 2005 |  | 2006 |  | 2007 |  | 2008 |  | 2009 |  | 2010 |  |
| --- | --- | --- | --- | --- | --- | --- | --- | --- | --- | --- | --- | --- | --- | --- |
| Test phase | 1 | 2 | 3 | 4 | 5 | 6 | 7 | 8 | 9 | 10 | 11 | 12 | 13 | 14 |
| **Herd A** |  |  |  |  |  |  |  |  |  |  |  |  |  |  |
| Total positive cows <sup>a</sup> | 14 | 25 | 34 | 28 | 34 | 32 | 21 | 23 | 24 | 17 | 14 | 9 | 7 | 0 |
| Animals tested | 315 | 330 | 364 | 349 | 354 | 364 | 338 | 332 | 337 | 347 | 341 | 347 | 296 | 239 |
| Apparent prevalence | 4.4 | 7.6 | 9.3 | 8.0 | 9.6 | 8.8 | 6.2 | 6.9 | 7.1 | 4.9 | 4.1 | 2.6 | 2.4 | 0.0 |
| New cases<sup>b</sup> | 14 | 16 | 18 | 12 | 13 | 13 | 7 | 10 | 11 | 6 | 5 | 2 | 0 | 0 |
| Cow-years at risk<sup>c</sup> |  | 239 |  | 293 |  | 284 |  | 272 |  | 276 |  | 280 |  | 198 |
| Apparent incidence<sup>d</sup> |  | 0.13 |  | 0.10 |  | 0.09 |  | 0.06 |  | 0.06 |  | 0.02 |  | 0 |
| **Herd B** |  |  |  |  |  |  |  |  |  |  |  |  |  |  |
| Total positive cows | 9 | 8 | 6 | 3 | 6 | 5 | 4 | 3 | 3 | 2 | 5 | 5 | 1 | 0 |
| Animals tested | 106 | 122 | 128 | 128 | 113 | 113 | 115 | 114 | 111 | 109 | 113 | 109 | 82 | 1 |
| Apparent prevalence | 8.5 | 6.6 | 4.7 | 2.3 | 5.3 | 4.4 | 3.5 | 2.6 | 2.7 | 1.8 | 4.4 | 4.6 | 1.2 | 0.0 |
| New cases | 9 | 1 | 2 | 0 | 5 | 4 | 1 | 0 | 1 | 1 | 4 | 1 | 0 | 0 |
| Cow-years at risk |  | 72 |  | 99 |  | 95 |  | 94 |  | 93 |  | 83 |  | 37 |
| Apparent incidence |  | 0.14 |  | 0.02 |  | 0.10 |  | 0.01 |  | 0.02 |  | 0.06 |  | 0 |

| **Herd C** |  |  |  |  |  |  |  |  |  |  |  |  |  |  |
| --- | --- | --- | --- | --- | --- | --- | --- | --- | --- | --- | --- | --- | --- | --- |
| Total positive cows | 0 | 17 | 26 | 23 | 19 | 22 | 18 | 20 | 18 | 15 | 13 | 8 | 7 | 0 |
| Animals tested | 0 | 121 | 145 | 149 | 178 | 161 | 145 | 155 | 157 | 145 | 142 | 117 | 102 | 0 |
| Apparent prevalence | NA | 14.0 | 17.9 | 15.4 | 10.7 | 13.7 | 12.4 | 12.9 | 11.5 | 10.3 | 9.2 | 6.8 | 6.9 | NA |
| New cases | 0 | 17 | 9 | 7 | 5 | 9 | 5 | 4 | 4 | 5 | 2 | 2 | 1 | 0 |
| Cow-years at risk |  | 13 |  | 114 |  | 123 |  | 110 |  | 117 |  | 108 |  | 33 |
| Apparent incidence |  | 1.27 |  | 0.14 |  | 0.11 |  | 0.08 |  | 0.08 |  | 0.04 |  | 0.03 |

<sup>a</sup>Test positive cows by considering enzyme-linked immunosorbent assay (ELISA) testing, fecal testing and tissue testing.
<sup>b</sup>Number of cows tested positive for the first time
<sup>c</sup>Observation time (in years) from entry in the study (at the first testing) until each cow tested positive or left the study (by culling, i.e. the infection status of cow is right censored)
<sup>d</sup>New cases per year / cow-years at risk

Fig3 shows the model predicted prevalence with a 95% confidence interval while fitting against the observed prevalence. It should be noted that our model confidence interval overpredicts the prevalence of herd B, but for other two herds it forecasts the better fitting. Through this model fitting exercise, our aim was to estimate the critical infection parameters for each herd, so that we can suggest herd specific intervention strategies.

**Fig 3.**
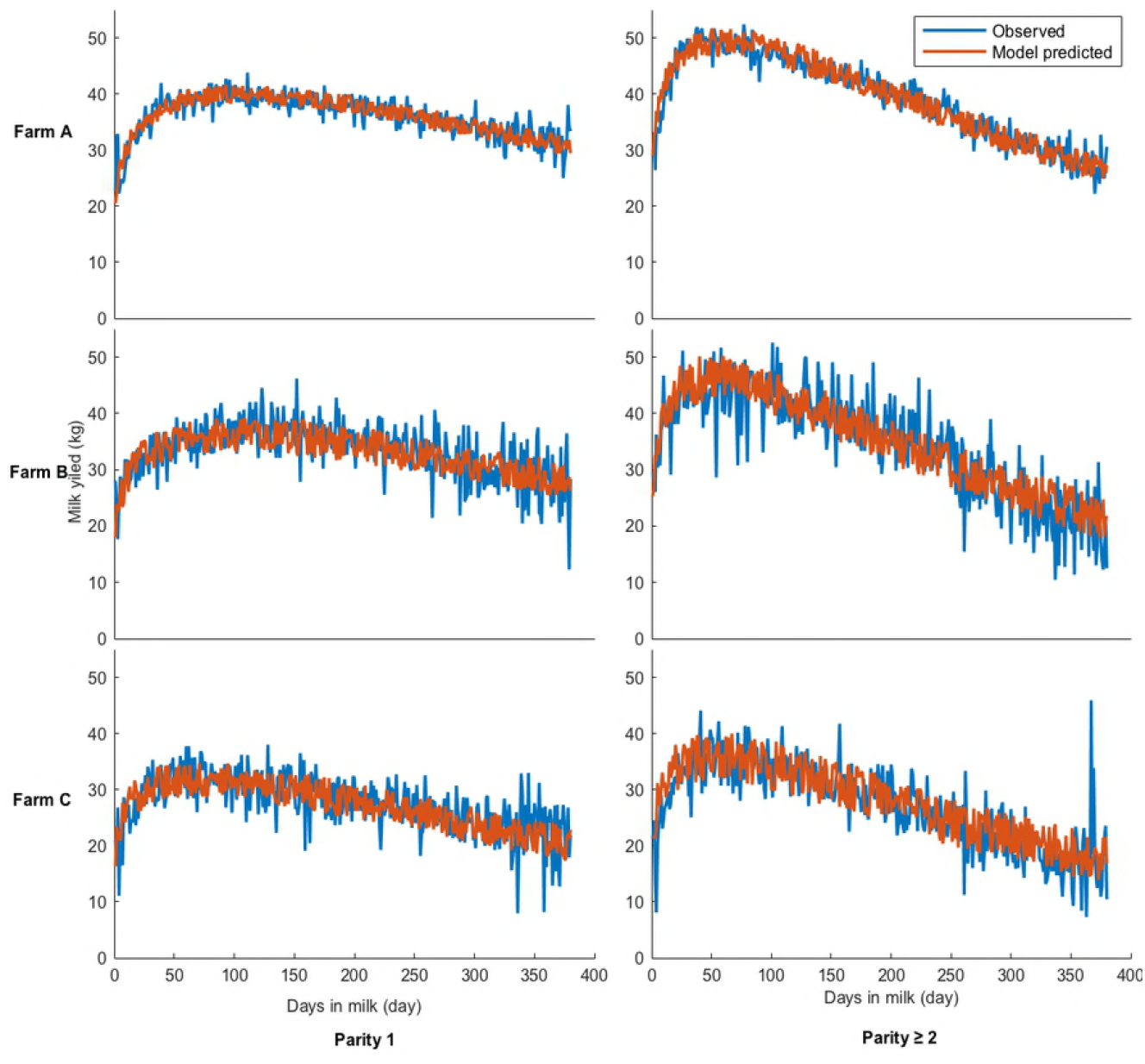
The fitting results of three *in silico* herds A (top), B (middle), and C (bottom) compared to the observed apparent prevalence for 7 years by biannual sampling. The shaded region shows the 95% confidence interval of the best 1% simulation runs.

### Estimated parameters

Table 5 provides the best fit estimates of herd-specific infection parameters for three northeastern dairy herds. Among the three herds, the model suggested that dam-to daughter transmission routes were the major transmission routes with the coefficient (β_*a*_) values of 0.4046, 0.1781 and 0.825 for farm A, B and C respectively. Environmental contamination was the second major transmission routes while adult-to-adult transmission route was ranked third. Interestingly, we found that the importance of adult-to-calf transmission was highest in herd C, in which the initial number of latent animals were highest in numbers among the three herds. Based on the best 1% parameter sets, herd C again had the highest number of latent animals present (shown in supplementary table S1).

**Table 5.** The values of fitted parameters for three farms A, B and C.

| Parameters | Herd A | Herd B | Herd C |
| --- | --- | --- | --- |
| Adult to adult transmission coefficient ($\beta_A$) | 0.0069 | 0.0023 | 0.0005 |
| Adult to calf transmission coefficient ($\beta_a$) | 0.4046 | 0.1781 | 0.825 |
| Environmental transmission coefficient ($\beta_{environment}$) | 0.0869 | 0.0711 | 0.0162 |
| Calf to calf transmission coefficient ($\beta_c$) | $5.3 \times 10^{-06}$ | $3.61 \times 10^{-06}$ | $5.2 \times 10^{-06}$ |
| Heifer to heifer transmission coefficient ($\beta_h$) | $4.36 \times 10^{-06}$ | $1.18 \times 10^{-06}$ | $1.98 \times 10^{-06}$ |
| Initial Latent animals ($H_i$) | 18 | 12 | 81 |
| Initial low shedding animals ($Y_{1i}$) | 15 | 2 | 12 |
| Initial high shedding animals ($Y_{2i}$) | 22 | 8 | 9 |

It is also noticeable that herd A has the highest adult-to-adult transmission probability among the three farms. Also, the initial starting distribution of the infected animals was very important for the fitting. It is seen that herd C start with the highest proportion of latent (73%) and low shedding (31%) animals among the three farms. The best-fitted parameters set is shown in the supplementary table (shown in supplementary table S1).

### Intervention strategies

Once the three *in silico* herds were obtained from the fitting exercises, our next aim was to test the risk-based test and culling policy for each farm. The risk-based intervention was implemented after 4 years of the initially fitted model to see the efficacy of the intervention strategy. Fig4 presents the summary of the pre-intervention, post-intervention and extended intervention results to the three fitted dairy herds. The results clearly show that the suggested intervention policy reduces the overall apparent prevalence for three herds, but it is noticeable that for high endemic herds the risk-based culling was comparatively less effective than the low endemic herds. To investigate further, we extended our intervention 2 years beyond the observations, but we did not see any elimination of MAP infection for the risk-based culling policy with control II. Culling red animals immediately (control I) was the best policy for all herds to decrease prevalence. Furthermore, we also calculated the number of years taken by the model to reduce the prevalence by 25% and 5% while two control programs were implemented after the pre-intervention period for three farms (shown in FigS2).

**Fig 4.**
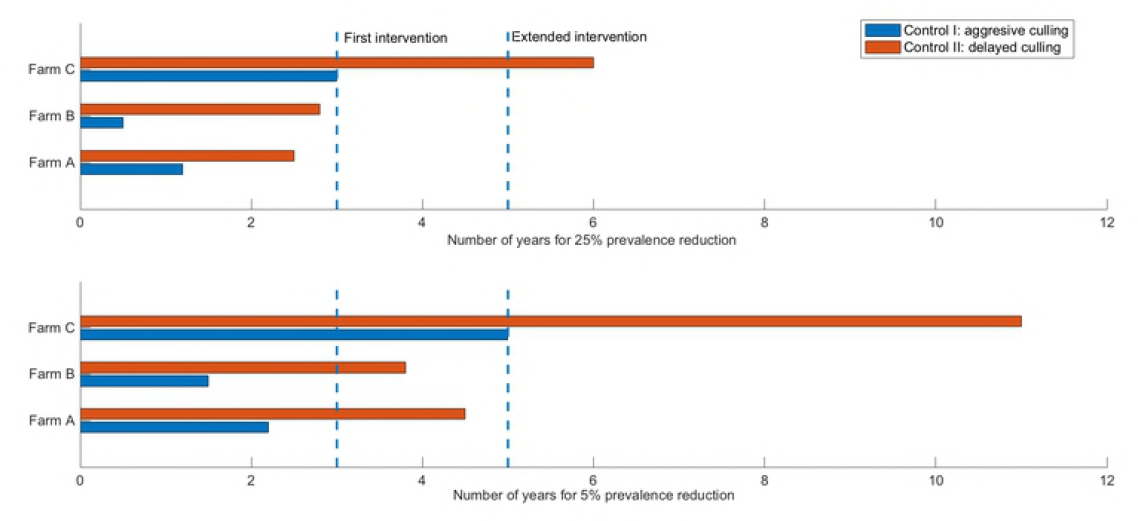
The apparent prevalence during the pre- and post-intervention period during the simulation of three *in silico* herds with two control strategies. Control I: culling red animals immediately and control II: culling only red animal with a delay of 305 days in milk. The two control measures are simulated in separating runs of the three *in silico* herds.

### Discussion and conclusion

Currently, only imperfect intervention strategies are available for PTB in the US. Therefore, there is a need to develop more effective control strategies to facilitate elimination of this disease from dairy herds. To enhance this effort, the mathematical modeling can play an important role, but the models can only provide realistic results when built from real herd data, to estimate the herd and infection-specific parameters and to test different intervention strategies prior to implementation in real herds. This paper presents an IBM modeling framework of MAP where simulation prediction was fitted and validated using datasets from a longitudinal study conducted in three northeastern dairy herds. The fitting exercise shows that the IBM is capable to reproduce the observed milk yield of each of the three herds separately and estimate key herd-related parameters. Next, the model results show the best fit to the observed apparent prevalence and estimate critical transmission parameters for three herds. Ultimately, the best fitted *in silico* herd models were simulated using risk-based test and culling intervention strategies, showing that these strategies may be more beneficial for low prevalence herds than for moderately endemic herds.

The epidemiology of MAP is difficult to study due to the slow progressing nature of MAP, insufficient testing methods, intermittent shedding of MAP and lack of clinical signs. Many infected animals are only detected years after initial infection or are actually never detected. However, precise information on the infection status of animals is valuable for implementing control strategies. Furthermore, specific information about the animal’s daily life events in the herd (such as age, milk yield, parity status, clinical signs and adult, calf and heifer rearing management policies) may assist in designing real-world control strategies. To this purpose, our IBM approach introduced a closed dairy herd model validated with longitudinal datasets (43,46,50). The basic herd fitting results suggest that we were able to create three *in silico* farms where the animal distribution was similar to the real herds (shown in table 1). This fitting exercise suggests that our base dairy herd model is capable of producing stable closed *in silico* dairy herds, with similar milk yield based on herd-specific milk yield parameters. This kind of features is very important to evaluate the economic efficacy of the implemented interventions (51). Moreover, often milk yield gets ignored from the MAP infection model, but accumulatively lower milk yield influences the culling of animals which is not normally marked that the animal was culled due to Paratuberculosis symptoms. Similar picture was seen in our data analysis of RDQMA herds where we found there were only 0.01% times where the animal was culled due to Paratuberculosis. Our previous study shows that low- and high-path animals produced more milk before their first positive test than always-negative animals, especially high-path animals. Although mean production decreased after a first positive test, low-path animals were shown to recover some productivity(50,52). To account the overall impact of milk yield on culling, we used threshold values of milk yield for parity 1 and 2 for each farm by calculating median milk yield values for each parity from observed data.

Next, we fitted three *in silico* herds to the apparent prevalence of the RDQMA herds. The 95% prediction interval shows that our model captured the trends of the apparent prevalence for three farms (shown in Fig3). Here we used antemortem ELISA and fecal testing and postmortem tissue and fecal testing results to determine the test positive animals in our model. In reality, determining the prevalence is a complex process and such fine-grained detail is rarely available. For antemorterm fecal culture test the sensitivity is determined 23-29% and 70-74% for infected cattle and infectious cattle respectively while at the slaughter house culture of tissue and fecal results 50% and 100% sensitivity and specificity, respectively. On the contrary for ELISA test our RDQMA suggests 20% and 96% sensitivity and specificity, respectively and these numbers are aligned with the previous reports by Nielsen and Toft(53). To avoid this complexity, we have chosen a range of 25-35% sensitivity for infected animals and 96% specificity. Recently, an adaptive test scheme was suggested from a simulation model simulated on the standard Danish dairy herd (8). In another study, test-records from 18,972 Danish dairy cows with MAP specific IgG antibodies on their final test-record were used to estimate age-specific sensitivities (54). It is a critical decision for a farm owner to choose one of the antemorterm test as the outcome of the fecal culture results can be delayed while ELISA test is also imperfect. Moreover, it also depends of the testing practices and recommendations varied in different geographical regions while strategies like adaptive test scheme, age-specific sensitivities and frequent testing can provide us optimal solution. But, care should be taken whether using frequent testing strategies may pick the false positive animal.

In order to control an infectious disease, it is important to determine which transmission routes are playing a major role in persistence of the pathogen on the farm. Traditionally, the dam-to-daughter route is considered the primary route for transmitting MAP, but it can vary due to herd management policy. It is very difficult to estimate this parameter directly from the epidemiological data due to imperfect testing, misidentification of super-shedders and management policies. The parameter value range estimated here suggests that dam-to-daughter transmission was indeed the primary transmission route with environmental transmission played as a secondary role. The role of environmental contamination is also difficult to measure from the epidemiological data, as MAP is pervasive within a dairy herd. A recent effort was made to quantify the environmental contamination through fecal-culture and mathematical studies (55). In our longitudinal data, the environmental samples were collected quarterly from several locations from farms. The cultures results suggest that manure storage areas and shared alleyways were most likely to be positive for three herds (56), but no relationship was found between non-pen environmental sample status and the distance between shedding animals and the sample’s location, and neighboring pens did not significantly affect the results of the pen-level analysis. In our model, we modeled β_*environment*_ in a crude way using a probability distribution for the sake of simplicity. To precisely quantify the role of different environments, further investigation into infection sources may be needed, potentially by examining the pathogens’ genomic sequencing data.

To date, the best-suggested control strategies against MAP is test and cull strategies. Previously, several compartmental models were used to test different testing and culling strategies, providing the average impact of the testing and culling strategies. However, targeted test and cull requires combining information from each individual animal with farm management and hygiene policy. Recently, an IBM model suggested that a new ethanol vortex ELISA (EVELISA) could be cost-effective and that quarterly test-and-cull control was able to significantly reduce the prevalence (41). Another model, SimHerd, developed by Kudahl et al. required fecal culture confirmation of ELISA-positive cows before culling, and relied on repeated testing to find the most infectious animal. Neither of these two models were validated and fitted to real dairy herd data (57). A recent mechanistic bio-economic model showed that MAP can be eradicated, although the control strategy necessary was economically unattractive (27,31). That model was parameterized specifically for Danish conditions, which are different from the US. In a previous effort, we suggested risk-based culling strategies with four different options: aggressive culling, culling open red cows after 305 DIM, culling dam and offspring and culling dam but not the offspring and we tested these intervention strategies along with different hygiene conditions on hypothetically endemic herds. For this study, we implemented two risk-based control strategies: aggressive culling and culling open red cows after 305 DIM on three pre-fitted herds. We found that aggressive culling resulted in the elimination of 24% and 47% of iterations after three years and extended intervention, respectively, for a very low endemic herd (farm B). We also found a probability of elimination 0.11 and 0.24 using culling of open red cows after 305 DIM in three years and extended intervention, respectively. However, it is expected to predict elimination in very low endemic herds and previously it was seen in a few studies (26,27,42,43,58). On the contrary, we found elimination in only 6% times after 5 years extended intervention using aggressive culling in case farm A, which has considerably higher prevalence, and we did not predict any elimination while culling open red cows after 305 DIM for farms C and A in long run. However, in terms of moderate and higher prevalence most of the cases, the farmers want to reduce the prevalence and it is important to simulate how long it takes to reduce the prevalence at a certain level. From FigS2, it can be said that low endemic herd is more likely to reach 5% of initial prevalence by less than 2 years while high endemic herd needs extended time to reach to that point, but it may take more than 10 years in some cases. This suggests that culling high shedding animals may not provide elimination in high endemic herds, although it can lower the prevalence. The study by Kirkby et al. serial testing along with hygiene play a critical role in the elimination process in Danish dairy herds, but these may not be economically justifiable (58). Caution should be taken in transferring conclusions from Denmark to the US, as both systems are different in many factors. Control activities are not uniformly coordinated nationally and internationally due to the variation in different farm management policies and government programs.

MAP is endemic in the bovine population in the US, which makes elimination unlikely at this time. When elimination is not possible, we have to rely on implementing the best herd-specific control strategies. Previous compartmental models have shown variable results for investigating infection dynamics(23,25,27), test-and-culling strategies(25,59), vaccination(24,60,61), and intermittent MAP shedding(30,43). None of these combined the individual animal’s information with herd management policy while fitting the model to real herd data, however. In this regard, the IBM paradigm should provide more effective approaches to test the intervention by considering information about the individual animal and overall population. Before using the insights of any IBM, very careful consideration should be given how the model was parameterized and validated. In this current study, we developed a fitting framework where an existing IBM model was fit against a longitudinal field study on three northeastern dairy herds to create the real herd’s condition in the *in silico* platform. The fitting exercises provide estimates of the critical parameters related to an infection whose transmission is herd-specific. Like all models, our model is limited by its assumptions. First, the current model fitting exercise only included combined testing efficacy, whereas in reality the observed herds used three different testing strategies (fecal culture, ELISA, and tissue culture). Second, the current model modeled the role of environmental contamination crudely, but the model is adaptive in nature, allowing for a more rigorous assessment of environmental contamination once data become available. Third, our current model did not include any economic justification of the suggested control strategies, but the same base model has previously been used to show the economic justification of culling in case of the MAP in a separate study (51).

This modeling and fitting exercise presented in this paper open multiple doors of further investigations in future. One extension of model can include the impact of MAP infection on milk yield while including the economics of milk production for these three farms. Previously, it shown that the mean milk production decreases after a first positive test, non-progressing animals were shown to recover milk productivity while progressing animals continue to exhibit a decrease in milk production, especially after their first high-positive fecal culture (52). This indicates there needs more investigation how to relate milk production loss as a function of MAP infection progression and testing results. Another extension of the model may include the clinical and molecular data of the infected animals. But adding molecular data will require more investigation how to find who infects whom parameters from the phylogenetic analysis (62,63). The current model is adaptive in nature to add strain specific data for each individual animal.

In conclusion, an important aspect of model building is to perform validation of the models to the real-life data. In this study, we developed an IBM framework for validating a dairy herd model and infection dynamics of the MAP to a longitudinal dataset. The assessment of model results leads us to the conclusion that the evaluation of model results is still a combination of intuitive model results, validation of the model with the quality data, assumptions that integrated into the modeling process and estimation of key critical parameters along with true biologics. This framework can be used in any infectious disease scenario to quantify the importance of key transmission routes, mapping individual-level data to population-level phenomena and decision making based on implemented intervention policies while considering between host transmission mechanisms within a closed population. In summary, the quality of the conclusions drawn from model studies is closely linked to the quality of the data used for estimation of the parameters and model validation. Models that have been validated with real-world data are more likely to produce useful and valid results.

## Acknowledgments

The authors gratefully acknowledge funding provided by the National Institute of Food and Agriculture of the United States Department of Agriculture through NIFA Award No. 2014-67015-2240.

## Author Contributions

Conceived and designed the experiments: MAM, Performed the experiments: MAM, Analyzed the data: MAM, RLS, and AN, Wrote the paper: MAM, Editing and reviewing: RLS, AN, and YTG. and YHS.

---

**Table S1.** The best 1% parameter sets were ranked from the parameter searching space.

| Parameters | Herd A<br>Mean (95% CI) | Herd B<br>Mean (95% CI) | Herd C<br>Mean (95% CI) |
| --- | --- | --- | --- |
| Adult to adult transmission coefficient ($\beta_A$) | **0.0033 (0.00062-0.0069)** | **0.0046 (0.0017-0.0075)** | **0.0041 (0.00047-0.0065)** |
| Adult to calf transmission coefficient ($\beta_a$) | 0.54 (0.11-0.96) | 0.37 (0.064-0.079) | 0.63 (0.055-0.9) |
| Environmental transmission coefficient ($\beta_{environment}$) | **0.053 (0.0089-0.090)** | **0.05 (0.0056-0.078)** | **0.046 (0.0036-0.087)** |
| Calf to calf transmission coefficient ($\beta_c$) | $0.69 \times 10^{-6}$ ($0.4 \times 10^{-6}$-$1.2 \times 10^{-5}$) | $6.5 \times 10^{-6}$ ($0.69 \times 10^{-6}$-$1.2 \times 10^{-5}$) | $8.4 \times 10^{-6}$ ($0.18 \times 10^{-6}$-$1.2 \times 10^{-6}$) |
| Heifer to heifer transmission coefficient ($\beta_h$) | $0.54 \times 10^{-6}$ ($0.53 \times 10^{-6}$-$0.1 \times 10^{-4}$) | $4.9 \times 10^{-6}$ ($0.29 \times 10^{-6}$-$0.11 \times 10^{-5}$) | $3.9 \times 10^{-6}$ ($0.35 \times 10^{-6}$-$0.1 \times 10^{-4}$) |
| Initial latent | 30 (3-75) | 10 (5-19) | 73 (52-85) |
| Initial low shedding animals | 18 (4-35) | 12 (2-36) | 31 (4-49) |
| Initial high shedding animals | 16 (7-23) | 11 (2-22) | 13 (2-23) |

**Fig S1.**
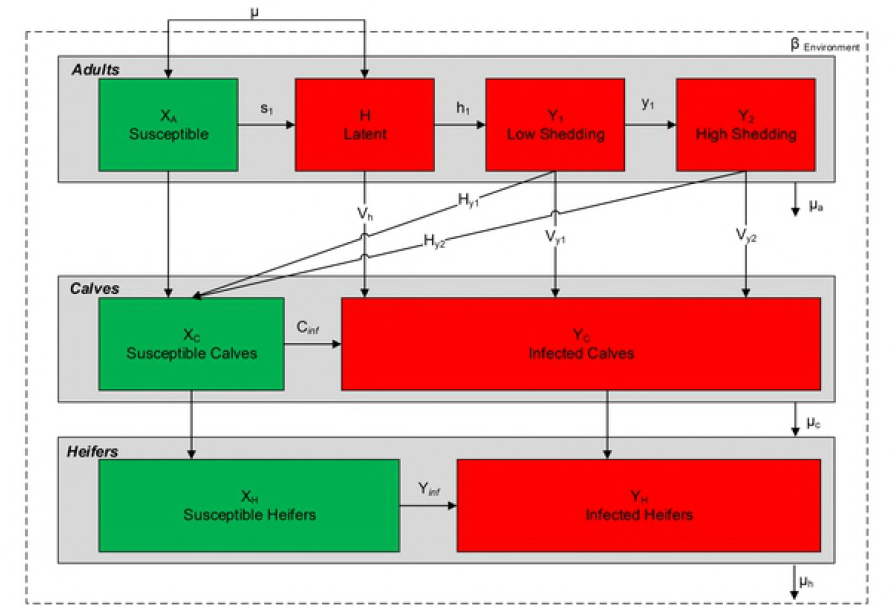
The model predicted fitted to the observed milk yield for 360 days in milk for Farm A, B and C. The milk yield was calculated using equation shown in the method section.

**Fig S2.**
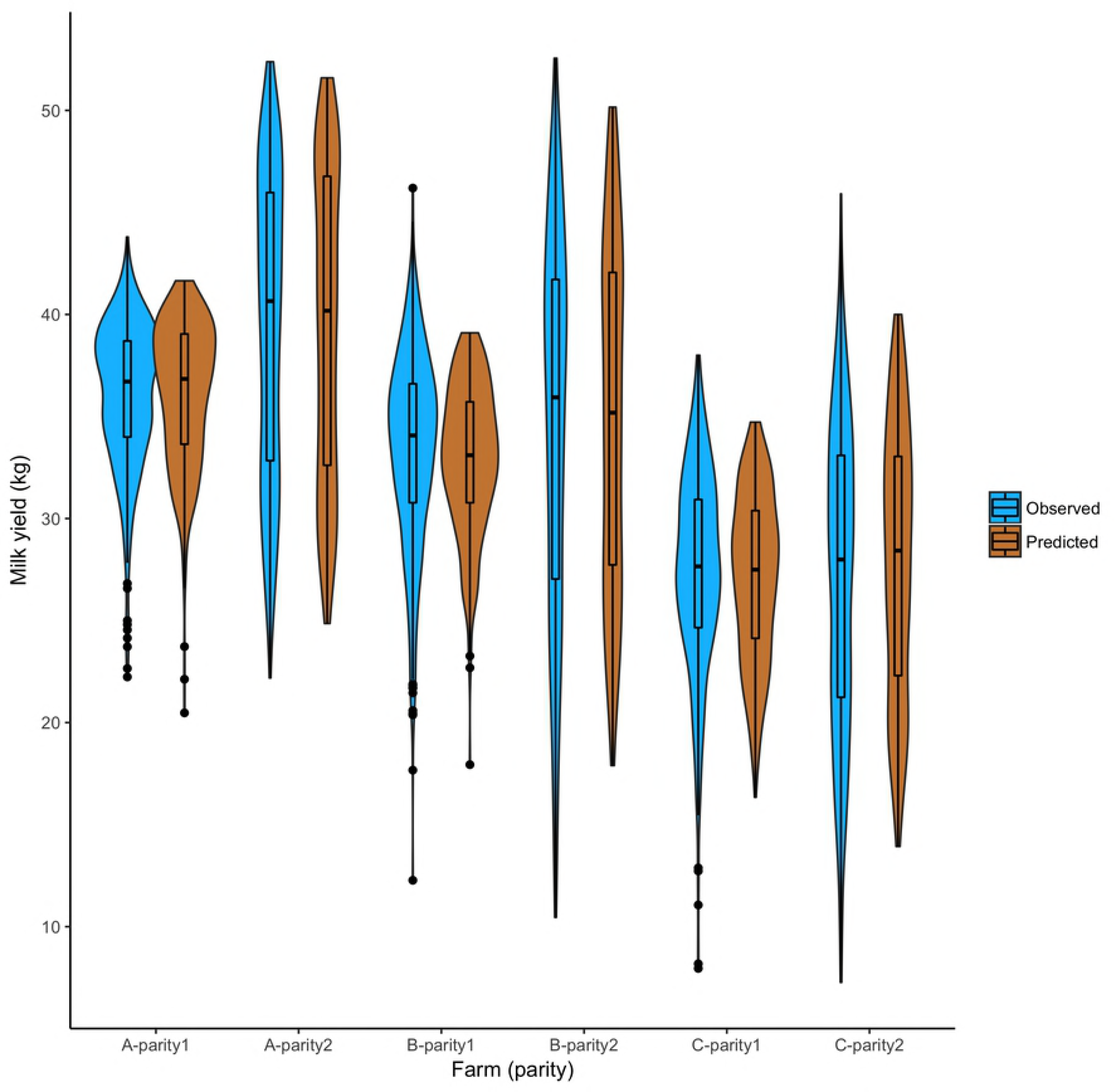
The model predicted median number of years to reduce the apparent prevalence by 25% (top panel) and 5% (bottom panel) calculated from top 1% simulations with best set of parameters while implementing two control scheme I: aggressive culling and control II: delayed culling after the pre-intervention fit for the farms A, B and C.

